# Archaeal self-activating GPN-loop GTPases involve a lock-switch-rock mechanism for GTP hydrolysis

**DOI:** 10.1101/2023.04.08.536109

**Authors:** Lukas Korf, Xing Ye, Marian S. Vogt, Wieland Steinchen, Mohamed Watad, Maxime Tourte, Shamphavi Sivabalasarma, Sonja-Verena Albers, Lars-Oliver Essen

**Author notes:** Contribution of both authors is considered equal (Lukas Korf, Xing Ye). Author Contributions: L.K., X.Y., M.S.V., S.V.A. and L.-O.E. designed research; L.K. and X.Y. performed experiments; W.S. performed and analyzed HDX-MS experiments; L.K., M.W. and L.-O.E. analyzed proteome data; L.K., X.Y., M.S.V. and L.-O.E. analyzed data; and L.K., X.Y., M.W., S.V.A. and L.-O.E. wrote the paper. **Competing Interest Statement:** The authors declare no competing interests.

## Abstract

Three GPN-loop GTPases, GPN1-GPN3, are central to the maturation and trafficking of eukaryotic RNA polymerase II. This GTPase family is widely represented in archaea but typically occurs as single paralogs. Structural analysis of the GTP- and GDP-bound states of the *Sulfolobus acidocaldarius* GPN enzyme (*Sa*GPN) showed that this central GPN-loop GTPase adopts two distinct quaternary structures. In the GTP-bound form the γ-phosphate induces a tensed dimeric arrangement by interacting with the GPN region that is relaxed upon hydrolysis to GDP. Consequently, a rocking-like motion of the two protomers causes a major allosteric structural change towards the roof-like helices. Using a lock-switch-rock (LSR) mechanism, homo- and heterodimeric GPN-like GTPases are locked in the GTP-bound state and undergo large conformational changes upon GTP hydrolysis. A *ΔsaGPN* strain of *S. acidocaldarius* was characterized by impaired motility and major changes in the proteome underscoring its functional relevance for *S. acidocaldarius in vivo*.

**Significance Statement:** GPN-loop GTPases have been found to be crucial for eukaryotic RNA polymerase II assembly and nuclear trafficking. Despite their ubiquitous occurrence in eukaryotes and archaea the mechanism by which these self-activating GTPases mediate their function is unknown. Our study on an archaeal representative from *Sulfolobus acidocaldarius* showed that these dimeric GTPases undergo large-scale conformational changes upon GTP hydrolysis, which can be summarized as a lock-switch-rock mechanism. The observed requirement of *Sa*GPN for motility appears to be due to its large footprint on the archaeal proteome.

## Introduction

GTPases are a large family of GTP-binding and-hydrolyzing enzymes that are widely distributed across all three domains of life ^1^. They contain a highly conserved GTPase domain (G domain) housing five fingerprint motifs responsible for coordination of GTP and catalysis: G1 (also P-loop or Walker A motif) interacts with the 5’ phosphate moieties of GTP, the G2 and G3 motifs are required for coordination of a magnesium ion essential for catalysis, the latter of which furthermore specifically accommodates the 5’ γ-phosphate of GTP, and G4 and G5 establish specific binding of the nucleotides’ guanine base ^2, 3^. As GTPases cycle between their GTP and GDP-bound states via their intrinsic GTP hydrolytic activity they often function as molecular switches differentially regulating a plethora of downstream effector proteins involved in crucial cellular processes ^2, 4, 5^.

Most GTPases belong either to the TRAFAC (Translation Factor Association) class involved in translation, signal transduction and intracellular transport, or the SIMIBI (Signal Recognition Particle, MinD and BioD) class; members of the latter engage in protein localization and trafficking, membrane transport and chromosome partitioning ^1, 4, 6^. Many well-studied GTPases, such as Ras and heterotrimeric G proteins ^7–9^, belong to the TRAFAC class, whose members generally do not form nucleotide-dependent dimers ^1^. In contrast, homo- and heterodimerization of SIMIBI class proteins like the signal recognition particle (SRP) and SRP receptor GTPases depends on ATP-or GTP-binding ^4,10^ and is hence relevant to regulation of their intrinsic ATP/GTP hydrolysis activity, specific interactions with effector proteins, and their biological functions ^6,^11^, 12^.

A novel type of SIMIBI GTPase, the GPN-loop GTPase, was discovered by structural analysis of the protein PAB0955 from the euryarchaeon *Pyrococcus abyssi* ^13^. PAB0955 adopts a homodimeric state, whose quaternary structure was found to be independent of GTP binding. This characteristic of GPN-loop GTPases sets them aside from other SIMIBI GTPases, which require switch-dependent changes of their quaternary state to fulfil their biological function. Another feature is the eponymous GPN motif in the G domain that is highly conserved in archaeal PAB0955 and eukaryotic XAB1-like orthologs. This motif (Gly-Pro-Asn) is inserted between the SIMIBI class motifs G2 and G3. GPN-loop GTPases were described to occur only in archaea and eukaryotes, but not in bacteria. Eukaryotes typically feature three GPN-loop GTPase paralogs: GPN1 (annotated before as Npa3, XAB1 or MBD*in*), GPN2 and GPN3. These paralogs play essential roles in nuclear localization and biogenesis of the RNA polymerase II ^14–17^. In yeast, deletion of genes encoding the XAB1 homologs GPN1, GPN2 or GPN3 was found to be lethal ^18^. Furthermore, yeast GPNs assemble into heterodimers with different yGPN combinations, which appear to be irrespective of the nucleotide-bound state of the GPNs as pull-down assays, FRET and molecular modeling studies indicate ^14, 19^. The biological function of archaeal GPN-loop GTPases remains unknown, although they exhibit the highest sequence similarities with eukaryotic GPN1 orthologs ^1, 19^. Nevertheless, the crystal structures of *Pyrococcus abyssi* GPN (PAB0955) and yeast GPN1 revealed a similar homodimeric topology, nucleotide coordination and a mode of catalysis, which involves the asparagine residue of the GPN motif ^13, 16^.

For long time the cellular function of GPN-loop GTPases remained enigmatic. *Homo sapiens* GPN1 was described first as XPA-binding protein 1 (XAB1) or MBD2-interacting protein (MBD*in*) because of its interaction with the nucleotide excision repair protein xeroderma pigmentosum group A protein (XPA) and methyl-CpG-binding protein MBD2, thus putting forward a role in DNA repair and methylation-dependent transcription ^20, 21^. Elucidation of the protein interaction network of all three GPN proteins in yeast revealed their essential role in the regulation of nuclear import of multiple RNA polymerase II (RNAPII) subunits ^22, 23^. The interaction of yeast GPN1/Npa3 with RNAPII subunit Rpb1 *in vitro* was stronger in presence of GTP than GDP ^24^. Furthermore, mutations in the G1, G2, G3 and GPN motifs of GPN1/Npa3 were either lethal or resulted in a slow growth phenotype in yeast ^23^, collectively suggesting a link between the nucleotide-bound state of GPN GTPases and their cellular function.

In the archaeon *Sulfolobus acidocaldarius*, the GTPase *Sa*ci1281 (from hereon: *Sa*GPN) represents the solitary GPN-loop GTPase. We previously identified *Sa*GPN as an interaction partner of the Ser/Thr specific protein phosphatase PP2A ^25^, which we anticipate to play an important role in intercellular signaling pathways because *S. acidocaldarius* possesses a diverse phosphoproteome ^26^. Interestingly, PP2A interacts with ArnA and ArnB, which are described as negative regulators of the assembly of the archaellum ^25, 27, 28^. PP2A deletion strains are therefore hypermotile and defective in cell size regulation and metabolism ^26^.

We investigated the *in vivo* function of *Sa*GPN in *S. acidocaldarius*, focusing on its potential role in the regulation of archaella synthesis. We also analyzed the protein structure and conformational changes of *Sa*GPN by crystallization and HDX analysis. The results from this study shed light on the *in vivo* function of GPN-loop GTPase in archaea, and we will also present a novel activation mechanism of GPN-loop GTPases.

## Results

### GPNs occur in all domains of life

To provide an overview of GPN-loop GTPases (IPR004130), we analyzed and visualized this protein family using a sequence similarity network (SSN), which provides insights into protein relationships and evolutionary origins based on different sequence alignment scores. Thereby, nodes correspond to protein sequences, and each edge represents one relation, which scales in stringency with decreasing E-values, allowing clustering based on this alignment threshold ^29^. Cluster analysis of IPR004130 with an E-value of 10^−40^ reveals that GPN-loop GTPases (GPNs) from archaea share a closer relation to GPN1 than to GPN2 and GPN3, which is consistent with the essential diversity of GPNs in eukaryotes ^15^. Moreover, GPN1 does not cluster with its sister paralogs GPN2/3, although the latter shares the same cluster, suggesting a greater functional discrepancy between GPN1 and GPN2/3 than between GPN2 and GPN3. Additionally, it appears that GPNs of TACK archaea, which include *S. acidocaldarius*, have a closer relationship with GPN1 than with other archaeal GPN-loop GTPases such as *Pyrococcus abyssi* (Figure 1). Furthermore, *Pyrococcus* and *Sulfolobus* GPNs begin to separate from each other at alignment scores with E-values of <10^−40^. Segregation became even more evident at a slightly increased E-value cutoff of <10^−45^, with the eukaryotic GPN1 clade being completely dissociated from archaeal GPNs. Likewise, *Pyrococcus*-like GPNs are almost completely clustered away from the *Sa*GPN-loop GTPases (Figure 1). This indicates, that despite sharing identity to some extent, these archaeal GTPases have some differences that are yet to be clarified. In our analysis of 1,541 unique archaeal sequences, belonging to 469 different organisms, we found that GPN-loop GTPases in archaea mostly occurred as singlets, with a total of 412 organisms. However, 57 archaeal organisms had two or more GPN paralogs, refuting the established assumption that archaea generally harbor only a single ortholog of GPN-loop GTPases ^16, 19, 30^. For bacteria, 4,331 sequences were used in the SSN, of which 1,859 distinct organisms were identified. Although GPN singlets also form the largest group in the bacterial domain with 762 organisms, the distribution among additional GPN loop GTPases is much more pronounced than in archaea. Accordingly, 410 organisms were identified with two GPN paralogs, and 386 with three. In addition, some bacterial species appear to have four or more GPN-loop GTPases, exceeding the number of three, GPN1-GPN3, known in eukaryotes. Nevertheless, bacterial GPNs share few relationships with either archaeal or eukaryotic dominated clusters and primarily cluster within their own, indicating that GPNs could play a different and more diverse role in bacteria than in eukaryotes or archaea. Furthermore, at an E-value of 10^−40^ archaeal orthologs assemble almost exclusively within a large cluster and share links with almost every other archaeal GPN, whereas bacteria split into several small clusters that have only minor relationships with the GPNs of their domain.

**Figure 1.**
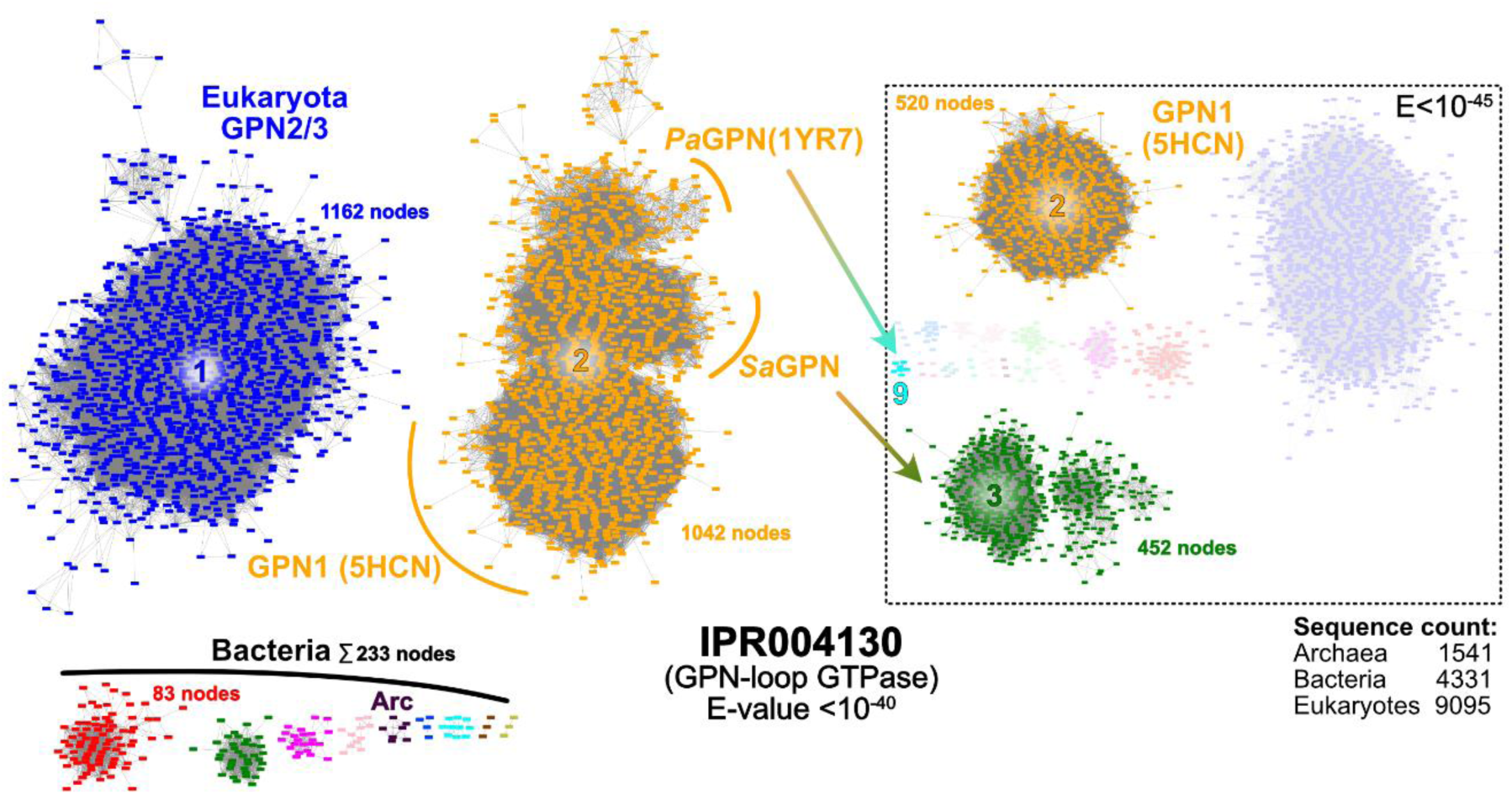
Sequence Similarity Network (SSN) of InterPro protein familiy IPR004130 (GPN-loop GTPases). The SSN shows conection between related proteins of the input protein familiy at the given E-value cutoff of 10^−40^. Cluster analysis numbers clusters based on member count, giving insights in overall cluster size. While the archaeal GPNs are clustered with GPN1 in cluster 2, other eukaryotic GPNs (GPN2/3) are located at cluster 1, revealing differences between GPN1 and its sister proteins. Additionally, cluster separation of cluster 2 is observeable, where *Pa*GPN, *Sa*GPN and GPN1 start to form their own subclusters. This is even more pronounced at higher stringency of 10^−45^ (dashed box), where *Sa*GPN separates from cluster 2 alongside most other archaeal GPNs into cluster 3 (452 nodes) and *Pa*GPN is clustered away into its own cluster 9 (cyan). Nodes not found in clusters 2 and 3 clustered in smaller/singleton subclusters, which are not displayed for clearity

### Biological effect of *saGPN* deletion on the archaeon *S. acidocaldarius*

*Sa*GPN was identified as a specific interactor of the serine/threonine protein phosphatase PP2A ^31^. In *S. acidocaldarius,* PP2A regulates growth, cell size and swimming motility ^26^. To investigate the physiological role of *Sa*GPN in *S. acidocaldarius*, a markerless in-frame deletion mutant was constructed. Deletion of *saGPN* did not affect cell growth (Figure 2A), whereas a pronounced decrease in swimming motility on semi-solid gelrite plates was observed (Figure 2B). However, electron microscopy (EM) analysis showed that Δ*saGPN* cells still assembled archaella, however, these samples have to be taken after 4 hrs of starvation as otherwise archaella cannot be observed even in the wild type (Figure 2C). To test whether deletion of *saGPN* affected *arlB* expression (encoding the archaellum filament protein, archaellin), Δ*saGPN* cells were starved for nutrients for 0–4 h, and *arlB* expression was analyzed by Western blot analysis and qRT-PCR (Figure 2D). *arlB* expression generally increased during nutrient starvation in both the wild type strain MW001 and Δ*saGPN* mutant. However, ArlB protein levels were significantly reduced in the Δ*saGPN* mutant compared with those in MW001. Similarly, decreased *arlB* expression was observed on the RNA level when comparing the Δ*saGPN* mutant with MW001 at 1, 1.5, 2, and 4 h after starvation (Figure 2E). Thus, *Sa*GPN affects the archaellum regulation network and apparently acts as a positive regulator in *S. acidocaldarius*.

**Figure 2:**
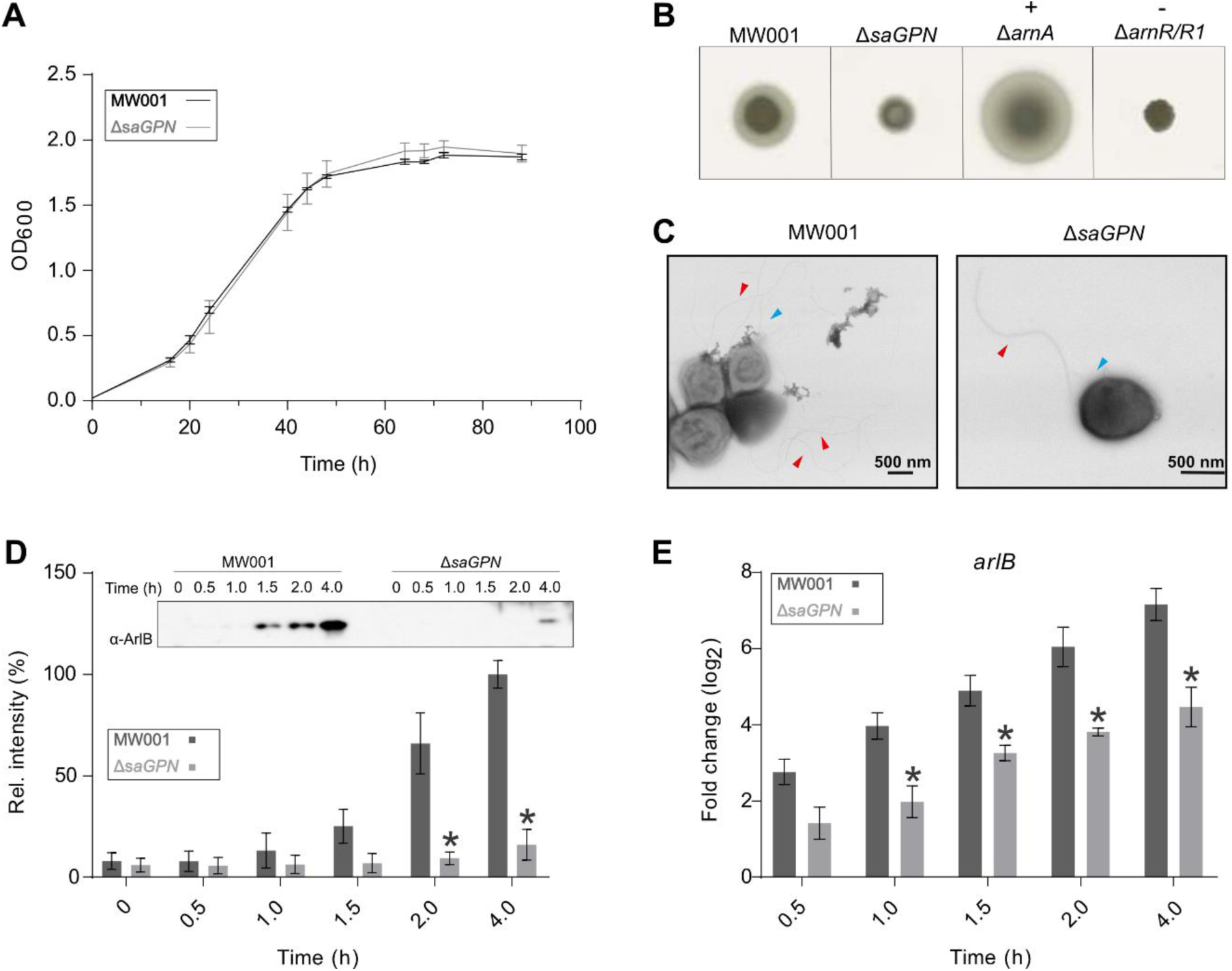
Deletion of *saGPN* impedes motility and ArlB (FlaB) expression in *S. acidocaldarius*. **(A)** Growth curves of *S. acidocaldarius* MW001 (black line) and Δ*saGPN* mutant (grey line) in nutrient rich medium. **(B)** Motility assays. Same amounts of cells from tested S. acidocaldarius strains were spotted on semi-solid gelrite plates and incubated at 75 °C for 5 days, then the plates were scanned and recorded. ΔarnA and ΔarnR/R1 strains were used as hyper-motile and non-motile control respectively. **(C)** Analysis of the archaellum formation in *S. acidocaldarius* strains. *S. acidocaldarius* MW001 and Δ*saGPN* knockout were cultivated in nutrient-depleted medium. After 4 h growth, cell samples were collected and applied to EM analysis. Red arrows highlight archaella and blue ones Aap pili. **(D)** ArlB expression on protein level and **(E)** arlB transcription level in *S. acidocaldarius*. *S. acidocaldarius* MW001 and Δ*saGPN* mutant were cultivated in nutrient-depleted medium for 4 h. Cell samples were taken at different time points, and analyzed by Western blotting with α-ArlB (left) and qRT-PCR (right). A representative Western blot is shown (D) and Western blots from biological triplicates were quantified. Relative transcription levels of arlB were normalized to secY. The values represent fold changes compared with the control from biological triplicates. Significant differences between MW001 (dark boxes) and Δ*saGPN* mutant (light boxes; p-value < 0.05) were indicated by an asterisk.

To explore whether *Sa*GPN affects other functions in *S. acidocaldarius* we performed a proteome-scale analysis of wild type and Δ*saGPN S. acidocaldarius* strains. These were grown as biological triplicates and samples were taken from nutrient-rich and nutrient-starved conditions. Each sample was measured as technical duplicates using a timsTOF (trapped ion mobility spectrometry, time of flight) mass spectrometer before averaging and further down-stream processing. Measurement of these proteomes led to the identification of 1,422–1,512 proteins per sample, which corresponded to a total of 1,627 different proteins of the 2,222 proteins encoded in the *S. acidocaldarius* genome. Comparing the two nutritional states of the WT, it is apparent that most of the proteome is nutrient-independent with only 37 proteins not identified in both states simultaneously (Figure 3A). For a quantitative assessment of the measured protein abundances data were further filtered using a *t*-test to analyze only hits, that showed statistically significant differences in abundance for data in their respective comparison. A total of 242 of the 1,492 (16.2%) overlapping proteins found in both nutritional states of the WT passed this filtering, indicating that expression levels of at least 16% of members of the proteome depend on nutrient availability. These data resemble previous label-based iTRAQ quantification analyses, which also indicated nutrient-dependent proteome changes of up to 12% ^32^. However, comparison of the Δ*saGPN* proteomes with their corresponding WT strains indicated that the amount of significantly altered proteins was approximately twice as high for the knockout strains, with 546/1,413 (38.6%) for nutrient-rich and 445/1,467 (30.3%) nutrient-starved conditions, respectively (Figure 3A). While this analysis reflected statistically significant changes, we also verified that the changes in protein levels were appropriately high by additionally filtering for proteins that showed a change of at least 50% in their respective comparison. This further filtering for largely altered expression levels (>50%) yielded 83 proteins for the WT_starved_/WT_rich_, 134 for the WT_rich_/Δ*saGPN*_rich_ and 127 for the WT_starved_/ Δ*saGPN*_starved_ comparison (Figure 3B). Heatmap analysis of this filtering shows that the distribution of over- and under-regulated proteins is similar between WT states whereas the knockout seems to have a more pronounced effect on downregulation. Nevertheless, the absence of the GPN-loop GTPase has profound effects on protein expression, even when only highly significant and major changes are considered.

**Figure 3.**
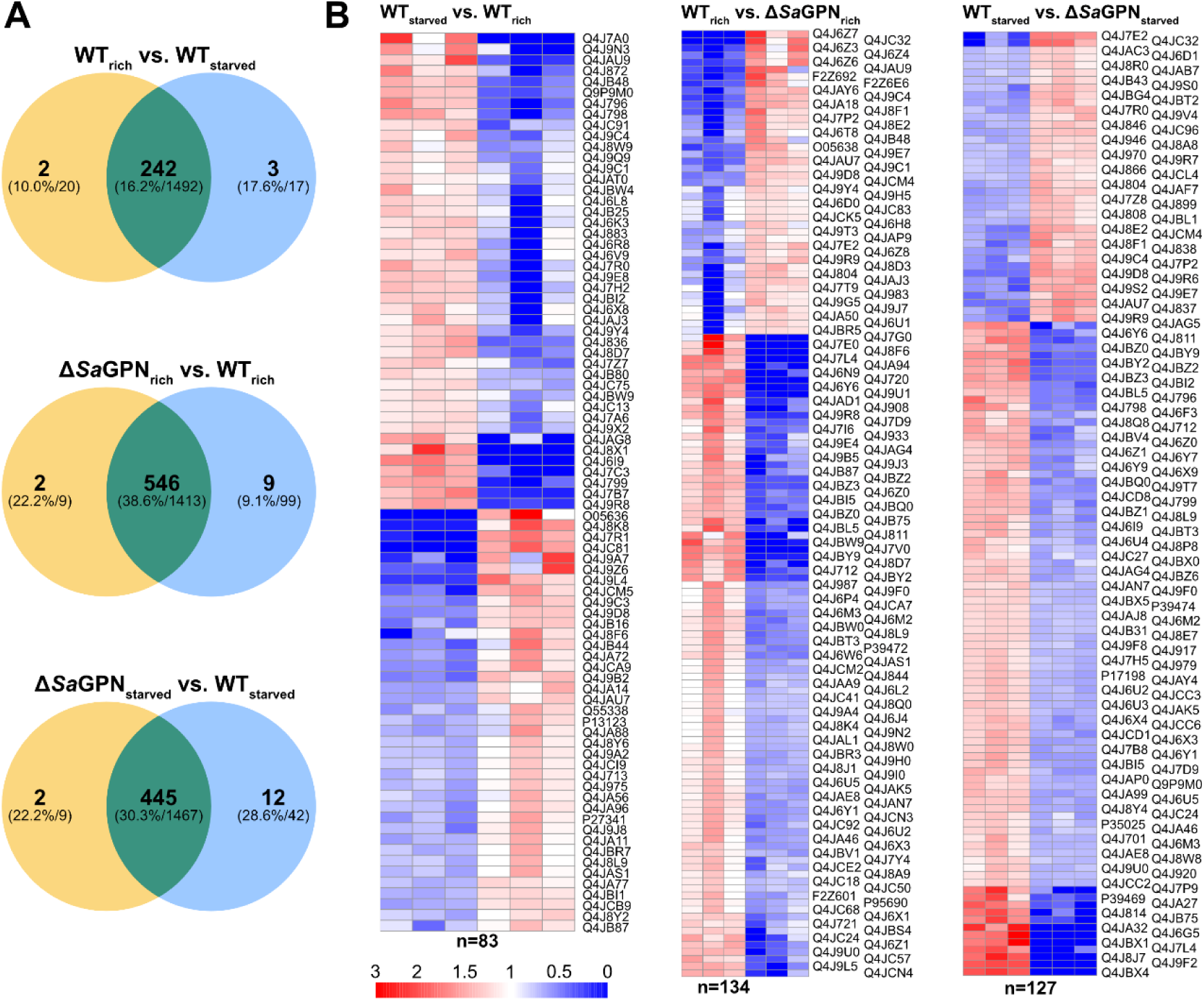
Knockout of *saGPN* has extensive ramifications on the *S. acidocaldarius* proteome. **(A)** V_ENN_ diagrams of proteomic data comparing different nutritional states of the wild type (WT) and *saGPN* deletion mutant. Diagrams show the number of overlapping and non-overlapping proteins that passed a one-paired *t*-test ≤0.01, to check for variation of the biological triplicates and the technical duplicates of their measurement, in bold. The proportion of these proteins filtered by the *t*-test to the total overlapping and non-overlapping proteins found is given in parentheses. The amounts of total proteins as identified by timsTOF, i.e. without *t*-test filtering, are given in absolute numbers next to them. **(B)** Heatmaps of proteins that exhibited highly affected expression levels (>50%) in their respective comparisons. The number of proteins that were severely affected is indicated below the maps. The scale bar represents the fold change of over-regulation (red) to under-regulation (blue). UniProt identifiers of the respective proteins are indicated next to the rows.

Evaluation of the Gene Ontology (GO) terms of the proteins shown in the heatmap analysis (Figure 3B) revealed that a large variety of different protein functions are affected by the *saGPN* knockout, pointing to a global role of *Sa*GPN in protein homeostasis (Suppl. Table 4). However, given the unaffected growth rate of the *saGPN* deletion strain, these changes in protein levels may correspond more to different lifestyles of this archaeal organism, e.g. due to the observed loss of motility (Figure 2).

### Biochemical and biophysical characterization of *Sa*GPN

In size-exclusion chromatography purified *Sa*GPN (30.5 kDa) eluted as a monodisperse peak at a volume corresponding to a molecular weight of ∼ 60 kDa, suggesting *Sa*GPN forms a dimer in solution (Figures S1A, S1B). GTPase activity of *Sa*GPN was highest in presence of Mg^2+^ at the optimal growth temperature (75 °C) of *S. acidocaldarius* (Figure S1C). However, this was not applicable for all experiments, as mechanical or chemical stress could result in an onset of protein aggregation at 75 °C, which is why the temperature was adjusted to 65 °C for kinetic experiments if not stated otherwise. *Sa*GPN preferred Mg^2+^ as the co-factor over other tested metal ions (Figure 4B). To determine the GTP hydrolysis kinetics, *Sa*GPN was incubated with 0-500 μM GTP at 65 °C in the presence of Mg^2+^. The calculated GTP hydrolysis was plotted against the GTP concentration and data were fitted with the Michaelis-Menten equation. The GTPase activity of *Sa*GPN had a K_M_ of 40.48 μM and a V_max_ of 6.4 nmol·mg^−1^·min^−1^ (36.4 nmol·L^−1^·s^−1^), respectively (Figure 4A).

**Figure 4.**
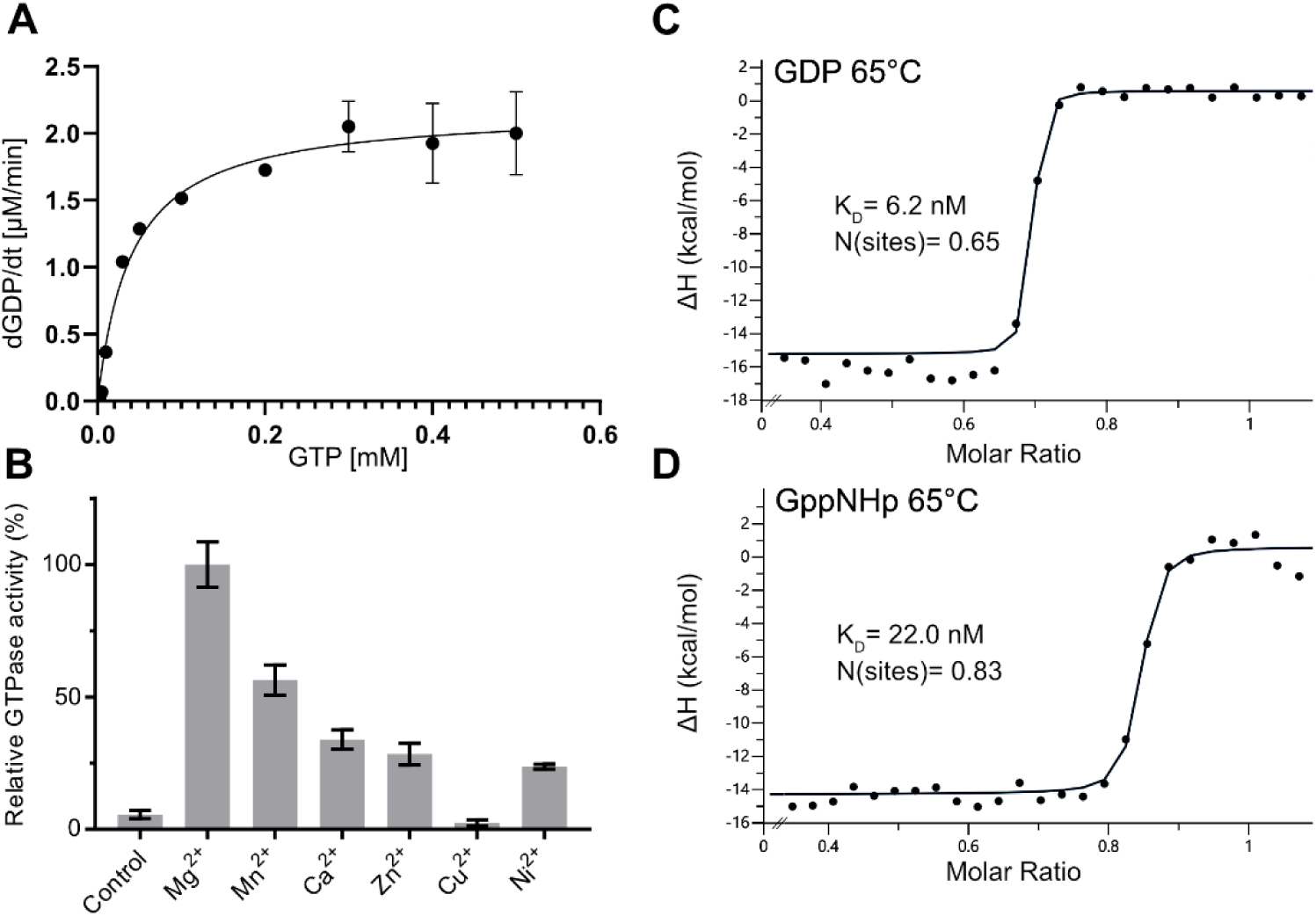
Despite showing low GTPase activity *Sa*GPNs nucleotide binding is very efficient. **(A)** M_ICHAELIS_-M_ENTEN_ Plot of *Sa*GPN activity at 65 °C, revealing an activity of 6.4 nmol hydrolyzed GTP per min and per mg of protein. Error bars (standard deviation) of some data points are too small to be visualized. **(B)** Effect of different bivalent metal ions on GTPase activity, which is highest with Mg^2+^. **(C)** ITC measurement of *Sa*GPN at 65 °C (to avoid premature protein degradation) with titration against GDP, showing a substrate affinity of KD = 6.15 nM for GDP. **(D)** Same measurement with titration against GppNHp, showing a substrate affinity of KD = 22.0 nM for GppNHp. Values are calculated from triplicate measurements.

Nucleotide binding efficiency of *Sa*GPN was investigated employing isothermal titration calorimetry (ITC), yielding dissociation constants (K_D_) in a low nanomolar range for all nucleotides investigated. To avoid protein degradation throughout the measurements, nucleotide affinity was determined at 65 °C, revealing K_D_ values of 6.15 nM and 20.03 nM for GDP and the non-hydrolysable GTP derivative GppNHp, respectively (Figures 4C, 4D). At 25 °C nucleotide affinity is even higher without significant disparity for triphosphate nucleotides, indicating that GppNHp is a suitable substitute for GTP (Figures S1D, S1E).

### *Sa*GPN overall structure

The crystal structure of *Sa*GPN was determined at a resolution of 1.8 Å (PDB: 7ZHF) and solved using molecular replacement, covering residues Y2-A240, which form ten α-helices and a five stranded parallel β-sheet, that is sandwiched by α5/α6 and α1/α10, respectively (Figure 5A).

**Figure 5.**
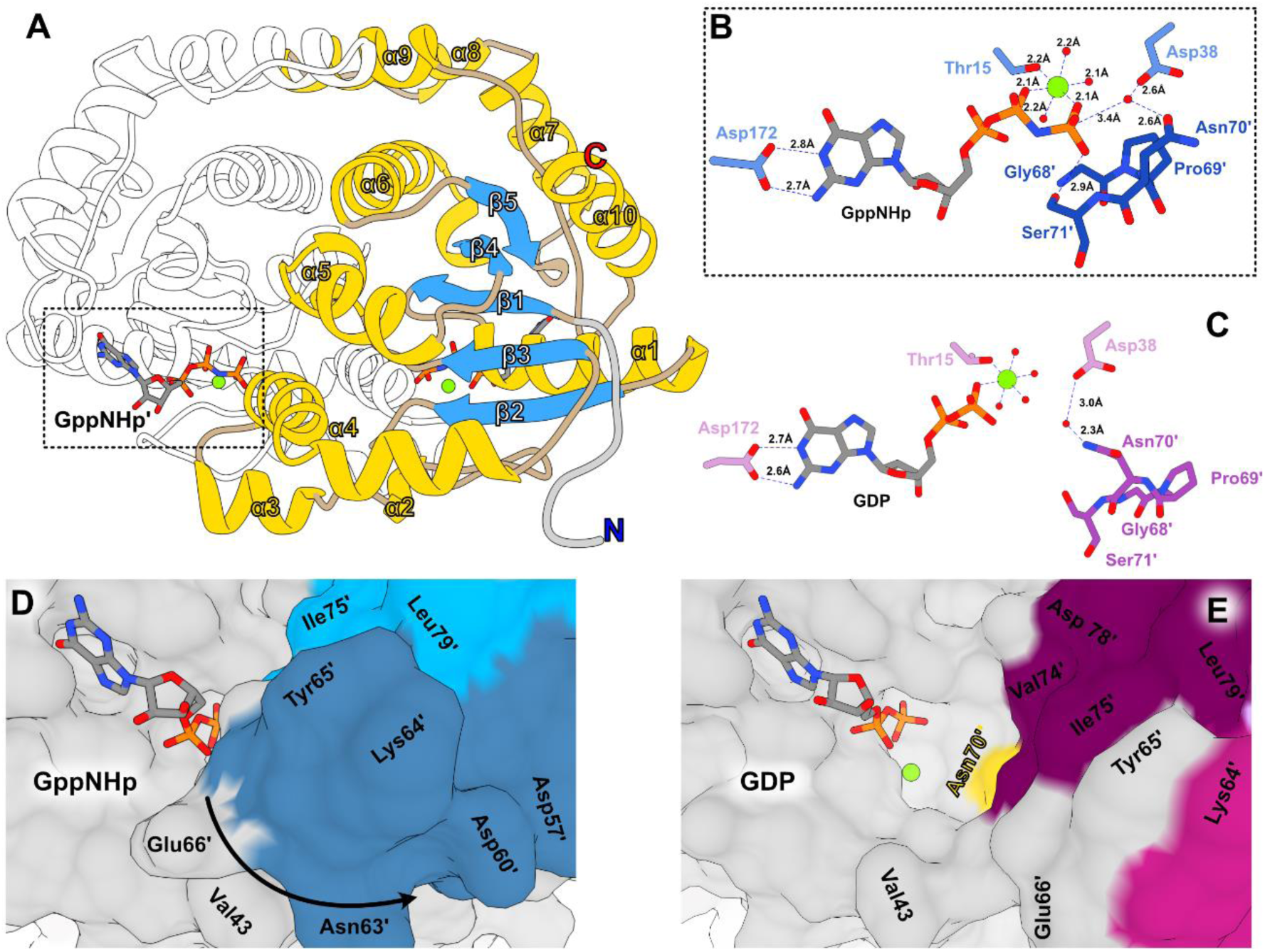
Overall structure of *Sa*GPN and substrate pocket. **(A)** Structure of *Sa*GPN with numbering of secondary structure elements on one protomer. Helices are colored in yellow, β-sheets in blue, disordered regions in wheat and the His-tag in grey. **(B)** Coordination of GppNHp in the substrate pocket with most important side chain interactions, including the near perfect Mg^2+^ octahedron and the catalytic active water coordination, with Mg^2+^ in green and waters in red. Primed residues refer to the second protomer. **(C)** Analogous coordination of GDP in the substrate pocket, revealing the difference in Mg^2+^ coordination as well as the increased distance towards the catalytic active water. **(D/E)** Surface view of the substrate pocket in its GppNHp (closed, PDB: 7ZHF) and its GDP state (open, PDB: 7ZHK) with coloring analogous to Figure 5A for better orientation.

Several small helices and helical turns consisting only of a few amino acids can be found throughout the whole structure including α2 and α8. Revealing only one protomer in the asymmetric unit, the crystal structure exhibits a homodimer for *Sa*GPN, concurring with the SEC elution volume (Figure S1). The dimer interface emerges alongside the surface of helices α4, α5, α6 and α9, whereby α6 contributes substantially to the dimeric assembly due to its core position allowing the formation of a hydrophobic pocket. This pocket is covered by a roof-like scaffold formed by α7-α9/α9’-α7’ laying orthogonally over α6 helices and parallel to the α2-α3 skid region, located at the opposite end. The eponymous GPN motif (residues 68-70) is found at the base of α4, which is involved in recruitment of the catalytically active water (N70) and substrate coordination at the substrate pocket of the second protomer. Nucleotide-binding is mainly guided by backbone coordination, however side chain interaction of D172 and K170 seems mandatory for coordination of the base and the ribose, respectively. Additionally, the Walker A motif (GxxxxGK[<u>T</u>/S], x=any amino acid) takes part in substrate binding, whereas K14 is responsible for the coordination and orientation of the phosphate section of the nucleotides, T15 is the only proteinaceous part of the octahedral Mg^2+^ envelope (Figure 4B) ^33^. Mg^2+^ coordination is also supported by the Walker B motif (hhhh<u>D</u>/E, h=hydrophobic), which interacts indirectly with the Mg^2+^ ion over a water intermediary via D102.

In addition to the GppNHp structure, we have succeeded in solving the *Sa*GPN structure in its GDP state (PDB: 7ZHK) at 2.4 Å resolution, allowing comparison between both states as well as with the only other known archaeal GPN-loop GTPase structure from *Pyrococcus abyssi* (*Pa*GPN). Apart from the missing γ-phosphate the binding mode of Mg^2+^•GDP to *Sa*GPN is comparable to that of Mg^2+^•GppNHp (Figure 5C). Furthermore, the distance between the catalytically active water is increased for D38 but only slightly shortened for N70’, suggesting that positioning of the catalytic water is driven by N70’ and activation by D38. However, due to the loss of the γ-phosphate interaction, the GPN loop detaches from the nucleotide, resulting in an open substrate pocket (Figure 5D, 5E). Therefore, the GDP state resembles a conformation that might exist prior to nucleotide exchange.

### Allosteric triggering of *Sa*GPN by bound nucleotides

While the difference of the quaternary structures of the GppNHp and GDP states is clearly apparent for *Sa*GPN (Figure 6), different nucleotide-bound states of *Pa*GPN only adopt an open state, which does not show significant changes upon nucleotide hydrolysis/exchange. Superposition of hydrolyzed and non-hydrolyzed states reveals this disparity, with r.m.s.d. values of 2.5 Å and 0.7 Å for *Sa*GPN and *Pa*GPN (GTP:1YR8/GDP:1YRB), respectively. The allosteric changes undergone by *Sa*GPN upon nucleotide hydrolysis correspond to global effects, which peak in a scissors-like movement of the α2-α3 skid region (Figure 6A). Achieving this scissors motion seems to involve many different local changes throughout the whole GTPase assembly. The roof helices shift against each other (Figure 6B) with the C-terminal α8 tails being pushed upwards, while α6 helices move towards and α7 drift away from each other. Moreover, α1 is stretched away from the skids and α4 bases move closer to each other, resulting in a convergence of the GPN motifs. In conclusion, a majority of the allosteric movement observable in the crystal structures takes place alongside the dimer interface, with most of the helical motion directed towards the roof structure. This includes α1 and α10, which are not part of the dimer interface yet contribute significantly to the scissors motion, with displacements of 18.9° and 12.0°, respectively (Figure 6C). As a result of the helical deflection toward the roof, β-sheet structures are elevated as well, releasing the tension needed to force the α2-α3 skids away, elongating the whole assembly.

**Figure 6.**
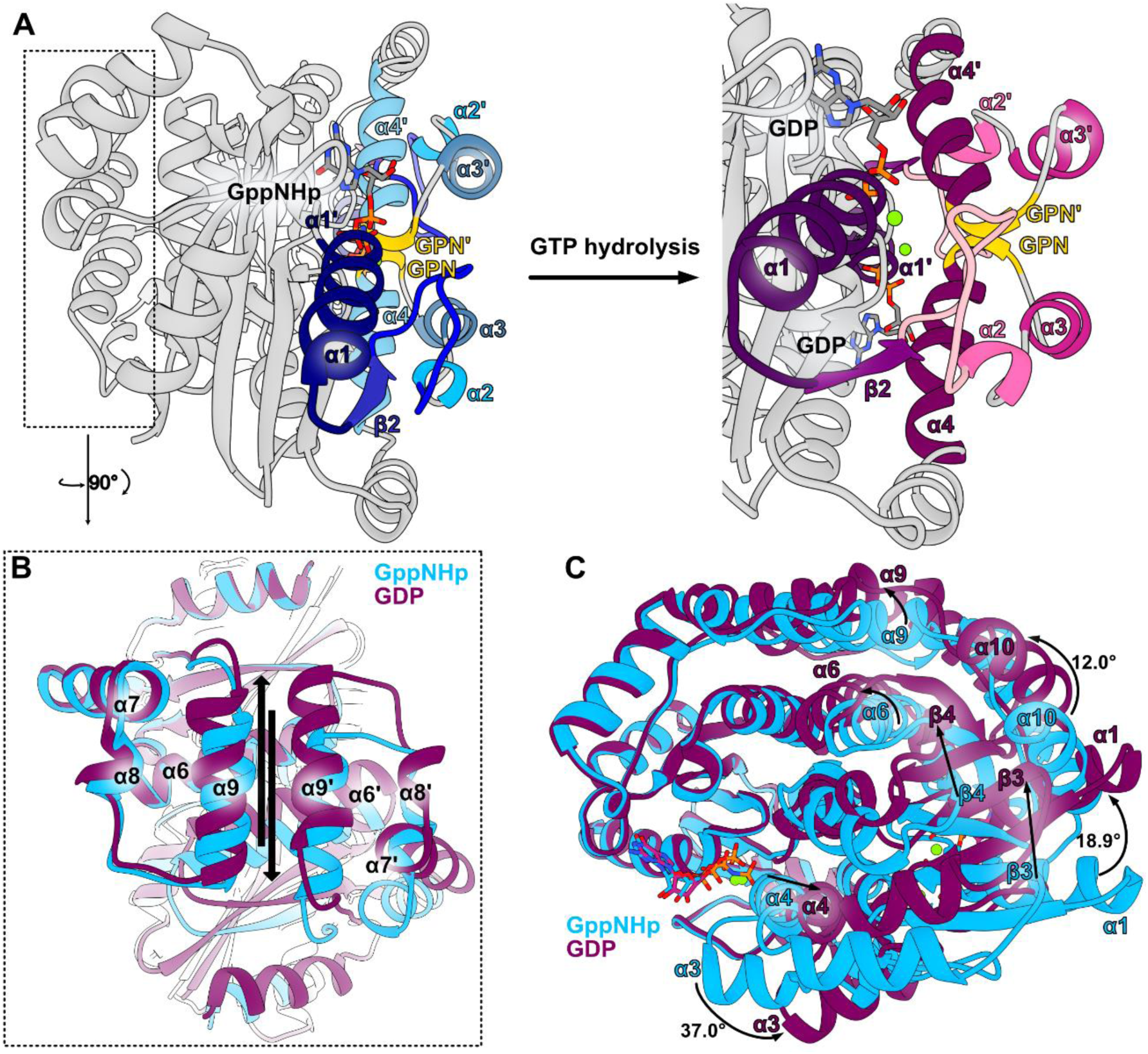
Allosteric changes of *Sa*GPN upon nucleotide hydrolysis observed in crystal structure. **(A)** *Sa*GPN skid region with corresponding allosteric changes upon nucleotide hydrolysis, revealing the push-out movement of the α3 skids. Secondary structure elements are colored in a blue gradient (GppNHp state) or purple gradient (GDP state) for better orientation and GPN motif in yellow. **(B)** Top down view on the roof helices with arrows indicating displacement between both states upon nucleotide hydrolysis. **(C)** Overall structure alignment of both nucleotide states, revealing major allosteric changes between states on outside helices as well as inside β-sheets. Most prominent helix shifts are given in degrees.

### In solution analysis of *Sa*GPN allosteric movements

We probed the nucleotide-dependent changes of *Sa*GPN in solution by hydrogen-deuterium exchange (HDX) coupled to mass spectrometry. Coordination of the GDP and GppNHp nucleotides by *Sa*GPN is evidenced from a reduction in HDX in the G1 motif (peptide 3-17), compared to the apo state (Figures 7A, 7B, S4). Further conformational changes induced by GDP encompass helices α3 and α4 (peptides 55-60 and 74-79, Figure 7B), which exhibit elevated HDX, and helix 8 and the subsequent strand β5 exhibiting reduced HDX (Figure S4). Binding of GppNHp to *Sa*GPN evokes even more pronounced perturbations in HDX (Figure S5). In addition to helices α3 and α4 incorporating more deuterium similar to the GDP-bound state, the roof-constituting helices α10-α12 specifically for GppNHp/*Sa*GPN incorporate more deuterium (peptides 151-168, 183-195 and 207-213, Figures 7A, 7B), which goes along with the vigorous movement of the roof helices observed in crystal structure. On the contrary, parts of helices α5 and α6 (peptide 111-121) close to the subunit interface incorporate less deuterium, collectively suggesting an altered topology of *Sa*GPN in presence of GppNHp. The disparate topology between GDP and GppNHp-bound *Sa*GPN is also reflected in a direct comparison of their HDX behaviors (Figure S6). However, while changes in HDX can be interpreted as changes in the structural assembly or movement of the region affected, the converse is not true.

**Figure 7.**
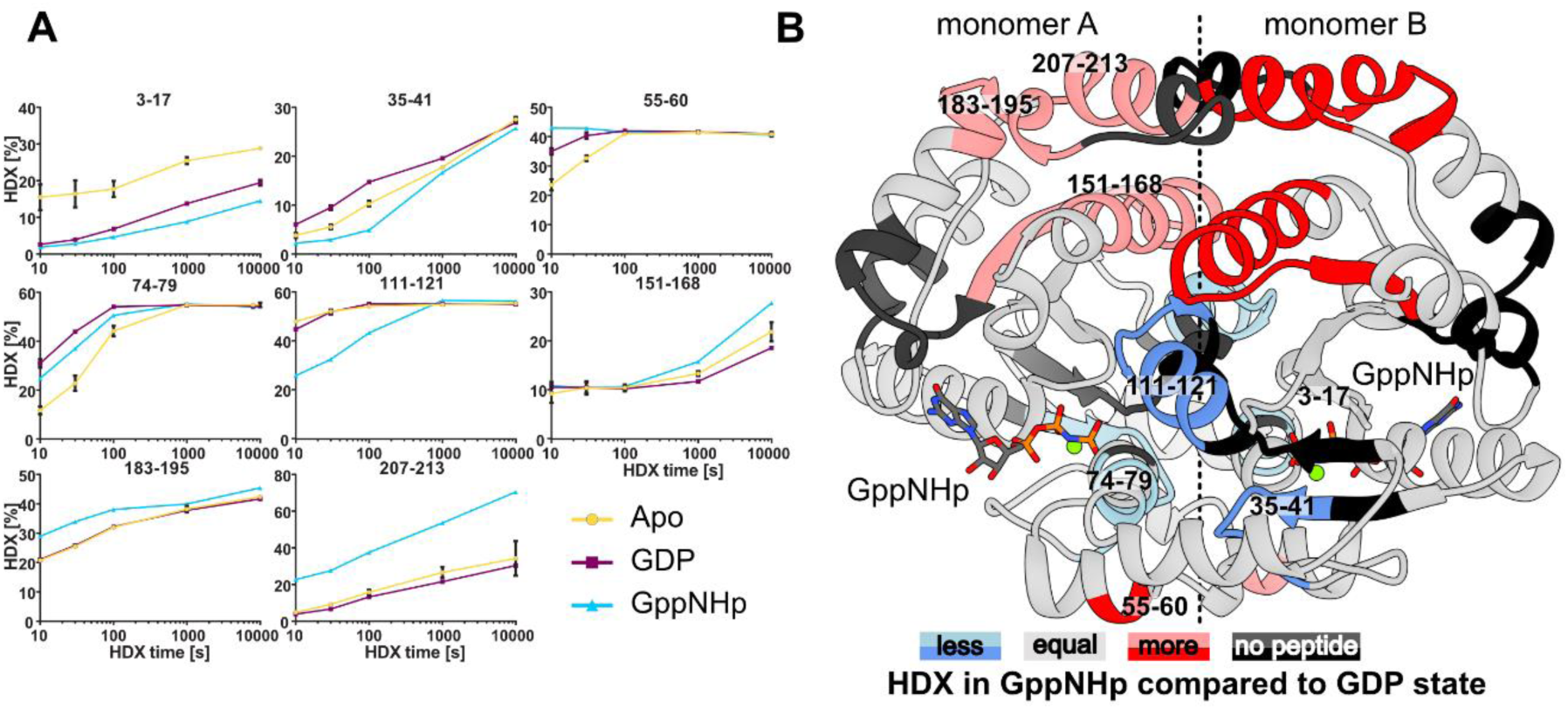
Allosteric changes of *Sa*GPN upon nucleotide hydrolysis observed in solution. **(A)** HDX of representative *Sa*GPN peptides covering regions of difference between the apo state (yellow circles), GDP-bound state (purple squares) and GppNHp-bound state (blue triangles). Data represent mean +/− s.d. of n=3 measurements. **(B)** Difference in HDX between GppNHp and GDP state mapped on *Sa*GPNs GppNHp structure (cumulative meaning that any HDX that happened of the respective region was mapped onto the structure with its highest difference). HDX data shows that differences between both states is most prominent in roof region, core elements and substrate coordinating/hydrolyzing regions, which goes along with most allosteric changes observed in the crystal structure. Numbers refer to amino acid position.

### GTPase activity and phosphorylation of *Sa*GPN are not required for motility

*Sa*GPN mutants (K14, D38A, D102A and GPN-AAA) were generated to investigate nucleotide binding and *in vivo* effects of defected mutants. All mutants were expressed in *E. coli*, purified and tested for their GTPase activity. GTPase activity was almost completely lost in all mutants (Figure S2A). When complementing the Δ*saGPN* strain with these *Sa*GPN mutants, all mutant proteins could be expressed in *S. acidocaldarius* (Figure S2C). Surprisingly, compared with the control, the swimming defect could partly be restored by these *Sa*GPN mutants, although they were catalytically inactive (Figure S7A). MANT-GTP binding assays indicated that these mutants still retained 20-50% of the wildtype GTP-binding capacity (Figure S7C), suggesting that nucleotide binding is unusually strong for a GTPase, since nucleotides generally fail to bind to G1/Walker A mutants ^34^.

Apparently, *Sa*GPN functions *in vivo* independently from its GTPase activity. Notably, a phosphoproteomics study of *S. acidocaldarius* demonstrated that *Sa*GPN can be phosphorylated at Y59 ^26^. A regulatory function for Y59 by (de)phosphorylation can be ruled as shown by the Y59F mutant. The *in vivo* and *in vitro* function of *Sa*GPN is unaffected by removal of this putative site of phosphorylation (Figure S7B).

## Discussion

GPNs are present in most organisms, in archaea occurring mostly as a single paralog and in eukaryotes in triple paralogs (GPN1-GPN3), where they perform non-redundant essential functions ^14^. However, little is known about this class of GTPases compared to other guanosine nucleoside phosphate hydrolyzing enzymes like small GTPases or G proteins. In 2007, GPNs were introduced by GRAS et al ^13^ as a self-activating, homodimeric GTPase family alongside the first GPN crystal structure from *Pyrococcus abyssi*. Our data now provide new insights into this understudied protein family of GPN loop GTPases.

We were able to solve the crystal structures for both a GTP trapped and a hydrolyzed nucleotide state of *Sa*GPN revealing major structural changes of the quaternary assembly for GPNs upon nucleotide hydrolysis, that have not been observed before. After GTP hydrolysis and loss of the γ-phosphate the substrate pocket switches from a closed into an open state by losing the interaction to the second protomer’s GPN motif. Opening of the catalytic cavity after nucleotide hydrolysis for subsequent nucleotide exchange goes along with strong nucleotide affinity of *Sa*GPN for both states. Nevertheless, self-activated GTPases like *Sa*GPN are capable of GTP turnover without the guidance of a guanosine exchange factor (GEF), at least *in vitro*. This is consistent of *Sa*GPN being a GTPase activated by dimerization (GAD), which are G proteins known for GTPase activity in the absence of GEFs or guanosine activation protein (GAP) ^3^. However, while dissociation constants of GADs to guanosine nucleotides are known to be within the low µM range, *Sa*GPN features an exceedingly high affinity with K_D_ values in the low nM range, matching the values of GTPases employing GEFs ^3, 10, 35, 36^. Furthermore, GPNs are dimeric regardless of their nucleotide-bound state, distinguishing them from most GADs ^10, 13^. Therefore, GPNs are different from most other G proteins and form a distinct class of their own. A comparably low GTPase activity appears to be a hallmark of archaeal GPN-loop GTPases as shown by turnover rates of 6.4 nmol/min/mg for *Sa*GPN and 12 nmol/min/mg for the orthologous *Pa*GPN ^3, 13^. These turnover rates may still allow a biological function for archaeal GPN-loop GTPases in the absence of GEFs.

However, due to the low intrinsic GTPase activity of GPNs both nucleotide states have a considerable half-life time, which is especially important for the GTP bound state, allowing for a potential regulatory function. Notably, eukaryotic GPNs are apparently not only involved in sister chromatid cohesion (human) and, in a chaperone-like manner, the assembly of the 12-subunit RNA polymerase II (*S. cerevisiae*), but also in other biological processes such as mitochondrial homeostasis and ribosome biogenesis ^14, 16, 37, 38^. Given this already broad range of functions for eukaryotic GPNs, archaeal GPNs with their largely GDP/GTP-affected quaternary structures may exert a similar range of functions, e.g. by acting likewise as a chaperone for multiprotein assembly.

Deletion of GPNs is known to be lethal in eukaryotes, but not in the case of the archaeon *S. acidocaldarius* indicating that archaeal and eukaryotic GPNs exert different functions ^13^. The Δ*saGPN* deletion mutant yields a phenotype with a highly diminished motility despite lacking any apparent effect on growth. This motility phenotype is independent from the catalytic capability of *Sa*GPN to hydrolyze GTP, as different G-box mutations, which cause lack of nucleotide binding and GTP hydrolysis, resulted in wild type-like motility. This finding suggests that the intrinsic GTPase activity of *Sa*GPN is dispensable for *S. acidocaldarius* motility behavior but leaves it open whether it may be of relevance to other processes not probed in this study.

Accordingly, a knockout of *saGPN* revealed that an unexpectedly large number of proteins with highly diverse range of functions are affected in their levels without causing a lack of viability for *S. acidocaldarius*. Compared to the metabolic adaptation of the *S. acidocaldarius* proteome upon starvation (Figure 3A), the even larger impact of the *saGPN* deletion suggests a key role of *Sa*GPN in protein homeostasis of *S. acidocaldarius*. Interestingly, the *saGPN* deletion also resulted in complete loss of an ortholog of the universal stress protein family, UspD (Uniprot: Q4JA32), which was undetectable in the proteome of the *saGPN* deletion strain, although it was highly abundant in all WT samples (Suppl. Table 4). UspD has very low sequence identity (8-11%) to known Usps of *S. acidocaldarius*, UspA-UspC, although it shares the typical Usp fold with the latter (Figure S8). Given that *Sa*GPN was found together with UspA in a previous co-IP assay ^25^, it is notable that UspA levels are almost unaffected in the *saGPN* deletion strain.

Since we were able to obtain snapshots of *Sa*GPNs catalytic cycle, we propose a mechanism, which we call ‘lock-switch-rock’ mechanism (Figure 8) and assign a catalytic function to the conserved Gly-Pro-Asn motif of GPN-loop GTPases. In addition to trapping a hydrolyzed and non-hydrolyzed nucleotide state of *Sa*GPN, it crystallized with its substrate pocket closed (PDB: 7ZHF), revealing the catalytic state immediately before hydrolysis, which is the first time that GPN-loop GTPases are observed in this state. This state builds up tension along the whole structure assembly induced by binding GTP, closing the GPN-lid and locking it in place. Accordingly, the distance between the nucleotide and the GPN loop increases upon nucleotide hydrolysis while the catalytic active water and the GPN’s asparagine (N70) retain their distance towards each other in both nucleotide-bound states. This suggests that N70 guides the positioning of the attacking water so it can then be activated by the catalytic aspartate of the G2 box (D38), followed by the nucleophilic attack of the hydroxide ion onto the γ-phosphate. The resulting hydrolysis to GDP and phosphate subsequently drives switching from a locked and tense state towards a relaxed state by performing a rocking motion effecting the whole structure.

**Figure 8.**
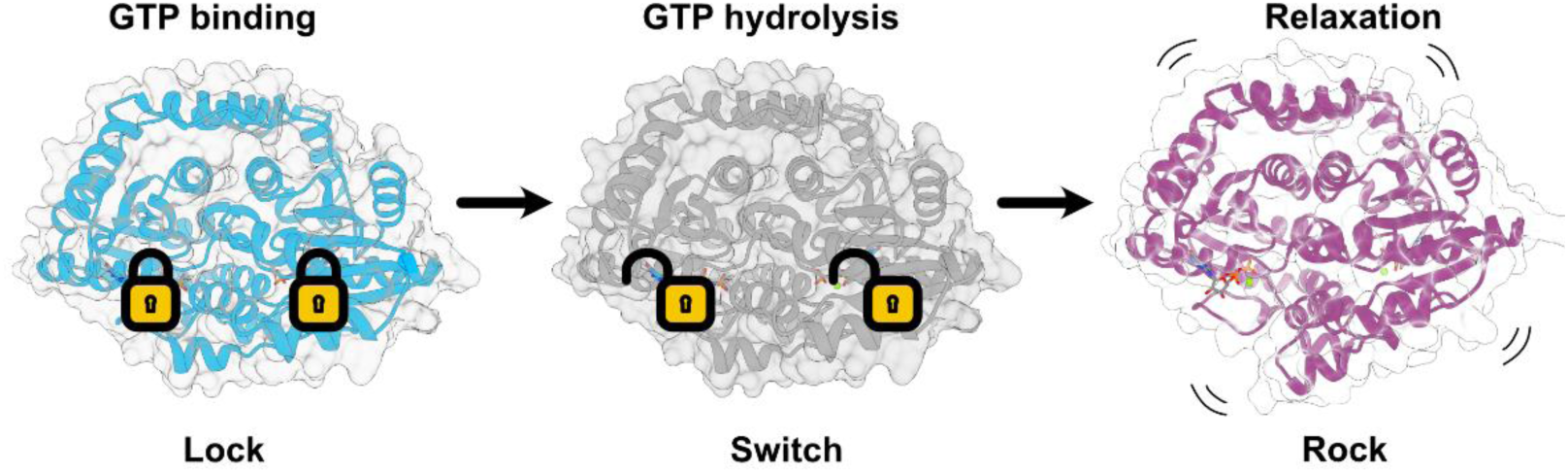
Lock-Switch-Rock (LSR) mechanism of *Sa*GPN. After GTP binding and before nucleotide hydrolysis, *Sa*GPN enters a locked state and stores tension throughout the structure. GTP hydrolysis leads to a loss of interaction in the GPN loop, which allows the transition from the locked state to a relaxed state in which a rocking motion is performed, stretching the GTPase assembly. Since these three steps represent the main catalytic states, we developed the lock-switch-rock mechanism, or LSR mechanism, for easy visualization and reference.

## Materials and Methods

### Strains and growth conditions

*S. acidocaldarius* wild type strain MW001 and all derived mutants in this study (Supplementary Table S1) were cultivated at 75 °C in Brock basal medium (pH 3.0-3.5) supplemented with 0.1% (w/v) NZ-amine, 0.2% (w/v) dextrin and 10 μg/mL uracil ^39, 40^. For *S. acidocaldarius* strains containing complementation plasmids, uracil was not needed.

### Construction of *S. acidocaldarius* Δ*saGPN* mutant

The plasmid pSVA5117 for the markerless in-frame deletion mutant Δ*saGPN* (Supplementary Table S1) was constructed as described previously ^40^. After transformation into *S. acidocaldarius* MW001, positive transformants were screened on 1st selection gelrite plates lacking uracil, then grown on 2nd selection gelrite plates with uracil and 5-FOA (100 μg/mL). Finally, Δ*saGPN* mutants were screened by colony PCR with checking primers (Supplementary Table S2), and further sequencing.

### Nutrient starvation assays and Western blots

Nutrient starvation assays and Western blots were performed as described ^31^. Briefly, at an OD_600_ of 0.4–0.5, overnight *S. acidocaldarius* cultures were collected at 75 °C and re-suspended in Brock medium in absence of NZ-amine and dextrin, followed by cultivation at 75 °C. *Sa*mples for the Western blot analysis were taken at the indicated times.

### RNA isolation and qRT-PCR

Total RNA samples were prepared with *S. acidocaldarius* cultures from nutrient starvation assays and qRT-PCR analysis was performed as described before ^25^. *secY* is a house-keeping gene, which was used as the reference gene for data normalization.

### Transmission electron microscopy

<u>Δ*saGPN*</u> and MW001 strains were grown in 50 mL Brock’s medium supplemented with 0.1% (w/v) NZ-amine, 0.2% (w/v) dextrin and 10 μg/mL uracil. At a OD600= 0.2-0.3 nutrient starvation was done for 4 hours to induce archaellation. Afterwards cells were applied on freshly glow-discharged carbon/formvar coated copper grids (300 mesh, Plano GmbH) and incubated for 30 seconds. This was repeated trice and excess liquid was blotted away. Cells were negatively stained with 2% Uranyl acetate. Imaging was done with Hitachi HT7800 operated at 100 kV, equipped with an EMSIS XAROSA 20 Megapixel CMOS camera.

### Motility Assays

Motility assays were performed as described ^40^. *S. acidocaldarius* strain Δ*arnA* and Δ*arnR/R1* were used as a hypermotile and non-motile control, respectively.

### PP2A phosphatase activity assays

Serine/threonine phosphatase activity assay of PP2A was performed using the artificial p-peptides RRA(pT)VA substrate (Promega) in lysis buffer containing 1 mM MnCl_2_ at 70 °C ^26^. The release of free phosphate from artificial p-peptides was measured using the Malachite green assay as described above ^41^.

### MANT-GTP Binding Assays

Binding of fluorescent MANT-GTP [(2’-(or-3’)-O-(N-methylanthraniloyl) guanosine 5’-triphosphate] to *Sa*GPN was measured by monitoring fluorescence increase upon binding to the protein utilizing a Fluoromax-4 spectrofluorometer (Horiba). The excitation wavelength was set to 285 nm, and emission wavelength was set to 450 nm. Slit widths for both excitation and emission were set to 10 nm. Binding was measured by incubation of 2.5 μM of *Sa*GPN/variants with MANT-GTP (5 μM) in buffer 50 mM MES pH 6.5, 150 mM NaCl, 5 mM MgCl_2_ at 25 °C. Fluorescence was corrected for MANT-GTP fluorescence in the absence of proteins. Fluorescence of *Sa*GPN wild type protein was set as 100% for relative MANT-GTP binding analysis.

### Sample preparation for proteomic mass spectrometry analysis

Strains were grown and subjected to 30 minutes starvation conditions as described above. Subsequently, 200 µL cell pellet equivalents were lysed in 900 μL urea buffer (8 M urea dissolved in 0.1 M NH_4_HCO_3_). After that, the samples were disrupted using glass beads and a FastPrep-24 MP Biomedical homogenizer (6.5 m/s, 3 cycles á 30 sec, 5 min resting on ice between cycles) with subsequent centrifugation (50 min, 14,000 rpm). Supernatant concentration was determined using Bradford assay and afterwards resuspended with the cell debris to yield full cell samples again. Protein concentration of full cell samples were estimated to be double the amount of the supernatant concentration. Samples were adjusted to 40 µL (8 M urea dissolved in 0.1 M NH_4_HCO_3_) containing 100-200 mg protein based on the concentration estimation and mixed with 1 µL TCEP (0.2 M) with subsequent incubation (1 h, 37 °C, 1,000 rpm). 1 µL iodoacetamide (0.4 M) was added and the solutions were incubated in the dark (30 min, 25 °C, 500 rpm) followed by addition of 1 µL *N*-acetyl-cysteine (0.5 M). Samples were incubated (10 min, 25 °C, 500 rpm) and diluted with 10.3 µL urea buffer (6 M urea dissolved in 0.1 M NH_4_HCO_3_) followed by the addition of 1.25 µL Lys C (0.2 mg/mL, 1:400 w/w) and digestion (4 h, 37 °C). Solutions were diluted with 145.3 µL urea buffer (1.6 M urea dissolved in 0.1 M NH_4_HCO_3_), mixed with 2 µL trypsin (1:100 w/w) and digested (overnight, 37 °C). pH was adjusted to <2 with 2.5 µL TFA (0.1% v/v). Samples were centrifuged (1 min, 14,000 rpm) and transferred to an equilibrated (0.1% v/v TFA) Chromabond C_18_ spin column (30 s, RT, 2,000 rpm). Peptides were eluted with 2x 150 µL elution solution (50% v/v ACN, 0.1% v/v TFA), dried in a vacuum centrifuge (45 °C, 4,000 rpm) and resuspended in 30 µL acetonitrile-TFA solution (10% v/v, 0.1% v/v). Samples were then measured using a timsTOF mass spectrometer in collaboration with the MarMass facility of Philipps University Marburg.

### Proteomics data evaluation

Data analysis of the timsTOF data was performed with MaxQuant 2.1.3 ^42^, sequence data for *S. acidocaldarius* DSM 639 (2,222 entries) were downloaded from Uniprot. For label-free quantification (LFQ) analysis, raw data were exported matching the quality and average area requirements described by the manufacturer. Protein abundances of biological replicates of the Δ*saGPN* strain and the wild type strain were filtered by a one-tailed, homoscedastic Student’s t-test (p < 0.01) using a mean abundance difference of > 10% as second criterion. The resulting lists of proteins were used for protein heat maps after filtering with a mean abundance difference of > 50% and visualization by the pheatmap R package. Gene ontology enrichment analysis was carried out using the GOATOOLS library, version 1.2.3, in python. In this gene ontology enrichment analysis, terms are represented for molecular function (MF), cellular component (CC) and biological processes (BP) of the identified proteins. To implement the GOATOOLS library, gene ontology terms (GO-Terms) for *S. acidocaldarius* DSM639 were downloaded using the QuickGO annotation online tool. A total of 9,487 annotations was available for *S. acidocaldarius* DSM639 proteins covering 1,548 gene products (70%); the remaining 674 are mostly still assigned as “conserved archaeal proteins”.

### Sequence Similarity Network (SSN) and *in silico* analysis

Generation of primary SSN data was done with the EFI-Enzyme Similarity Tool web service (UniProt Version: 2021_03; InterPro Version 87) based on InterPro Family IPR004130 (GPN-loop GTPases) with an UniRef90 restraint due to the size of the IPR ^29, 43^. SSNs were generated for E-values 10^−40^ and 10^−45^ with subsequent cluster analysis (standard options) and colorization. Analysis and visualization of the generated network was performed with Cytoscape 3.8.2 using the yfiles organic layout ^44^.

### Expression and Purification of *Sa*GPN

Recombinant overexpression of N-terminal His_6_ tagged *Sa*GPN was performed in an expression media containing 10 g/L tryptone, 10 g/L NaCl, 5 g/L yeast extract and 12.5 g/L lactose, employing BL21(DE3) Rosetta cells (37 °C, 150 rpm, 18 h). Harvesting (20 °C, 5,000 rpm, 20 min) was followed by resuspendation in lysis buffer (150 mM NaCl, 50 mM Tris, 15 mM EDTA, pH=8.0) and cell lysis via French Press. The lysate was centrifuged (18,000 rpm, 20 °C, 20 min), the supernatant heat treated (55 °C, 15 min) and centrifuged again (18,000 rpm, 20 °C, 20 min) before being loaded to a Ni-NTA column (5 mL). After sample application, the column was washed with 9 column volumes wash buffer (150 mM NaCl, 50 mM Tris, 25 mM imidazole, 15 mM EDTA, pH=8.0) and eluted with 5 column volumes elution buffer (150 mM NaCl, 50 mM Tris, 500 mM imidazole, pH=8.0). The elution was concentrated (<2.5 mL) and applied to a HiLoad^®^ 16/60 Superdex^®^ 200 pg (*GE Healthcare*) size-exclusion column, which had been equilibrated with running buffer (150 mM NaCl, 50 mM Tris, 10 mM MgCl_2_, pH=8.0). Fractions containing *Sa*GPN with a 260/280 nm ratio ≤0.55 were collected, concentrated, frozen in liquid N_2_ and stored at −80 °C.

### GTPase assays

GTPase activity of *Sa*GPN was measured using the Malachite green assay ^41^. For a 600 µL reaction mix at 65 °C 60µL10x*Sa*GPN (c_fin_=10 µM) was mixed with 60 µL 10xGTP stocks and adjusted to 600 µL with a buffer containing 50 mM Tris pH 8.0, 150 mM NaCl and 10 mM MgCl_2_. For each time point 60 µL of that mixture were taken and quenched with the Malachite green reaction solution. After incubation for 30 min the Malachite green solution developed its color and was measured. The absorption at 620 nm was measured and a phosphate standard was used for quantification.

### Crystallization of *Sa*GPN

Crystallization screens were performed as sitting drop experiments with 24.5 mg/mL *Sa*GPN and 3 mM nucleotide in running buffer mixed in a 1:1 ration with the respective crystallization condition, using 0.3 µL of each. Crystallization was observed in different conditions of all JCSG Core suits (*NeXtal*), however, the crystal used for the structure determination of *Sa*GPN(GppNHp) crystallized in the presence of 200 mM NaCl, 100 mM NaOAc (pH=4.6) and 30% (v/v) MPD (JCSG Core III/G11) after 24 h. *Sa*GPN(GDP) was crystallized in the presence of 200 mM MgCl_2_, 100 mM MES (pH 5.5) and 40% (v/v) PEG 400 (JCSG Core II/G3) after 24 h.

### Structure determination of *Sa*GPN

Data collection was performed with the Swiss Light Source at Paul Scherrer Institute in Switzerland. Phasing was done employing molecular replacement using BALBES (GppNHp state) and PHASER (GDP state) of the ccp4 pipeline followed by model building with PDB-REDO and ARP/wARP ^45–49^. Structure refinements were done by multiple rounds of manual model building with Coot followed by phenix.refine ^50, 51^. Structural analysis was carried out with PyMOL and visualization with Chimera 1.15 ^52, 53^.

### Isothermal titration calorimetry (ITC)

ITC experiments were carried out with purified *Sa*GPN concentrated to a 100 µM stock solution and stored at −80 °C as 300 µL aliquots. For each run, *Sa*GPN aliquots were thawed just prior to measurement, as were nucleotides (800 µM stocks). The sample cell of the MicroCal PEAQ-ITC, (Malvern) was heated to 65 °C or 25 °C, before injection of 40 µL syringe volume (nucleotides) over 52 injections under constant stirring. Evaluation and visualization of ITC data was performed with Malvern evaluation software.

### Hydrogen-deuterium exchange (HDX) mass spectrometry (MS)

HDX-MS experiments on *Sa*GPN were carried out similar as described previously ^54^. The samples contained 50 µM *Sa*GPN and 5 mM nucleotides (GDP, GppNHp) in a buffer containing 50 mM Tris-Cl pH 8.0, 150 mM NaCl and 10 mM MgCl_2_. Preparation of the HDX reactions was aided by a two-arm robotic autosampler (LEAP technologies). For deuterated samples, 7.5 μL of *Sa*GPN solution (with or without nucleotides) were supplemented with 67.5 μL of D_2_O-containing buffer to start the exchange reaction. After 10, 30, 100, 1,000 or 10,000 s at 25 °C, samples (55 µL) were taken from the reaction and mixed with 55 µL of quench buffer (400 mM KH_2_PO_4_/H_3_PO_4_, 2 M guanidine-HCl, pH 2.2) kept at 1 °C. 95 µL of the resulting mixture were injected into an ACQUITY UPLC M-Class System with HDX Technology (Waters, ^55^. Undeuterated samples of *Sa*GPN were prepared similarly by 10-fold dilution of the protein solution with H_2_O-containing buffer. *Sa*mples were flushed out of the loop (50 µL) with H_2_O + 0.1% (v/v) formic acid (flow rate of 100 µL/min) and guided to a column (2 mm x 2 cm) packed with immobilized porcine pepsin and kept at 12 °C for proteolytic digestion. The peptic peptides thus generated were collected on a trap column (2 mm x 2 cm) filled with POROS 20 R2 material (Thermo Scientific) kept at 0.5 °C. After 3 min, the trap column was placed in line with an ACQUITY UPLC BEH C18 1.7 μm 1.0 x 100 mm column (Waters), and the peptides were eluted at 0.5 °C column temperature using a gradient of H_2_O + 0.1% (v/v) formic acid (A) and acetonitrile + 0.1% (v/v) formic acid (B) at a flow rate of 30 μL/min as follows: 0-7 min/95-65% A, 7-8 min/65-15% A, 8-10 min/15% A, 10-11 min/5% A, 11-16 min/95% A. The peptides were ionized with an electrospray ionization source (250 °C capillary temperature, 3.0 kV spray voltage) and mass spectra acquired in positive ion mode over a range of 50 to 2,000 *m/z* on a G2-Si HDMS mass spectrometer with ion mobility separation (Waters), using Enhanced High Definition MS (HDMS^E^) or High Definition MS (HDMS) mode for undeuterated and deuterated samples, respectively ^56, 57^. Lock mass correction was implemented with [Glu1]-Fibrinopeptide B standard (Waters). During each chromatographic run, the pepsin column was washed three times with 80 µL of 4% (v/v) acetonitrile and 0.5 M guanidinium chloride, and blank injections were performed between each sample. All measurements were carried out in triplicate.

*Sa*GPN peptides were identified from the undeuterated samples with the software ProteinLynx Global SERVER 3.0.1 (PLGS, Waters), using the amino acid sequence of *Sa*GPN, porcine pepsin and their reverted sequences as database, and their deuterium incorporation determined with the software DynamX 3.0 (Waters) as described previously ^54^.

## Data availability

Protein structure data is deposited in the PDB and can be accessed via the codes 7zhf (*Sa*GPN GppNHp structure) and 7zhk (*Sa*GPN GDP structure). Remaining data are found in the supplementary material.

## Acknowledgments

We thank the beamline staff of the Swiss Light Source (SLS), PSI, Villigen, Switzerland, for support, the staff of Marburg crystallization facility (MarXtal) for technical support and Marta Rodriguez-Franco for assistance during EM imaging. We acknowledge support by the DFG through the Core Facility for Interactions, Dynamics and Assembly of biomolecular Structures (MIDAS). This work was supported by the Chinese Scholarship Council (PhD scholarship to X.Y.). L.-O.E. and S.V.A. thank the Life? program of the Volkswagen Foundation for funding. We would also like to thank the EM facility at the Faculty of Biology, University of Freiburg, for access to the TEM for generation of data. The TEM (Hitachi HT7800) was funded by the DFG grant (project number 426849454) and is operated by the University of Freiburg, Faculty of Biology, as a partner unit within the Microscopy and Image Analysis Platform (MIAP) and the Life Imaging Center (LIC), Freiburg.

